# Ambiguity-Aware Multi-Stage Cell-Type Annotation for Spatial Transcriptomics

**DOI:** 10.64898/2026.06.21.733596

**Authors:** Md Ishtyaq Mahmud, Veena Kochat, Humaira Anzum, Suresh Satpati, Jagan Mohan Reddy Dwarampudi, Kunal Rai, Tania Banerjee

## Abstract

Spatial transcriptomics enables characterization of cellular organization in intact tissue, but robust cell-type annotation remains challenging due to heterogeneous expression profiles, mixed populations, and transitional states. Existing methods often enforce a single label per cluster, obscuring biologically meaningful ambiguity and producing overconfident assignments.

We propose an ambiguity-aware, multi-stage framework for spatial cell-type annotation. The method combines hybrid spatial–feature clustering with constrained language-model inference over curated label sets, and assigns confidence scores based on marker coverage, candidate separation, and entropy. Low-confidence clusters are selectively refined via local reclustering of ambiguous regions, while unresolved clusters are preserved as mixed rather than forcibly labeled.

Applied to 10x Genomics Xenium spatial transcriptomics data from cholangiocarcinoma, the proposed refinement reduces cluster-level ambiguity from 16.1% to 2.27% and cell-level ambiguity from 18.4% to 0.86%, while improving confidence calibration. Spatial ablation confirms that topological integration resolves structural ambiguity over feature-only baselines, while constrained inference via a lightweight language model ensures scalable and biologically coherent annotations. These results highlight the importance of explicit ambiguity handling for reliable spatial annotation in heterogeneous tumors.

## 1 Introduction

Spatial transcriptomics enables high-resolution mapping of gene expression within intact tissue, providing insight into tumor heterogeneity [1]. In cholangiocarcinoma, spatially organized epithelial, stromal, and immune compartments often exhibit overlapping transcriptional programs and transitional states, complicating cluster-level annotation. Conventional pipelines rely on global clustering (e.g., Leiden or Louvain), while more recent approaches incorporate spatial constraints via hybrid spatial–feature graphs or graph neural networks to jointly model neighborhood structure and gene expression [2, 3, 4]. Although effective for capturing large-scale tissue organization, these methods may obscure fine-grained or mixed subpopulations within heterogeneous tumor microenvironments.

Recent work has further integrated large language models (LLMs) and foundation models for cell-type annotation [5, 6, 7]. While leveraging semantic priors improves consistency, these approaches often rely on high-capacity models and enforce single categorical assignments, even when transcriptional evidence is diffuse. Refinement-based strategies revisit uncertain cluster assignments [8, 9], but ambiguity is typically treated as noise rather than as a biologically meaningful signal.

We argue that ambiguity should be explicitly quantified and preserved when supported by evidence. We therefore propose an ambiguity-aware, multi-stage framework combining hybrid spatial–feature clustering with constrained language-model inference and confidence scoring. Low-confidence clusters are selectively refined via local reclustering of induced subgraphs, and unresolved clusters are abstained rather than forcibly labeled. We evaluate the framework on cholangiocarcinoma spatial transcriptomics data and assess robustness across language-model capacities.

The key contributions of this work are:

1. An ambiguity-aware annotation framework with confidence scoring based on coverage, margin, and entropy.
2. A multi-stage refinement strategy that selectively reclusters low-confidence clusters using restricted spatial–feature graphs.
3. Structured representation of mixed clusters through composite or state-level annotations.

## 2 Proposed Method

We propose an ambiguity-aware, multi-stage framework for spatial cell-type annotation that integrates hybrid spatial–feature clustering with constrained language-model inference and confidence-based abstention. The pipeline consists of: (i) hybrid graph construction and initial clustering, (ii) constrained semantic annotation with confidence scoring, (iii) selective multi-stage refinement of low-confidence clusters, and (iv) confidence-based abstention from definitive labeling when separability criteria are not met.

### 2.1 Hybrid Spatial–Feature Clustering

Given a spatial transcriptomics dataset with gene expression matrix *X* ∈ *R*^*n*×*p*^ and spatial coordinates for *n* spots or cells, we first compute a low-dimensional representation using hierarchical sparse non-negative matrix factorization. Let *Z* ∈ *R*^*n*×*k*^ denote the learned embedding.

We construct two adjacency graphs, namely,

1. Feature graph *A*_feature_: a *k*-nearest-neighbor graph computed in the hSNMF embedding space.
2. Spatial graph *A*_spatial_: a graph encoding physical adjacency based on spatial proximity. These graphs are combined into a hybrid adjacency matrix:

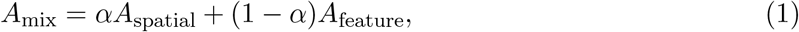

where *α* ∈ [0, 1] controls the contribution of spatial structure. Leiden clustering is then performed on *A*_mix_ to obtain initial cluster assignments. The mixed adjacency matrix is row-normalized prior to clustering.

### 2.2 Marker Extraction

For each cluster, differential expression analysis is performed against all other cells using the `scanpy.tl.rank_genes_groups` function (one-vs-rest comparison) in the Scanpy Python library. This procedure computes log_2_ fold changes, raw and adjusted *p*-values, and test statistics for each gene using the Wilcoxon rank-sum test. From these results, we additionally derive mean expression within and outside the cluster and expression prevalence. The top-ranked marker genes per cluster are retained for downstream semantic annotation.

### 2.3 Constrained Language-Model Annotation

To support constrained language-model annotation, we construct a curated candidate label set from PanglaoDB [10] and CellMarker [11]. Gene symbols were standardized to HGNC nomenclature and synonymous labels were harmonized, and unsupported marker–label mappings were removed. The resulting reference panel was used solely to constrain the admissible label space during semantic inference, with candidates filtered by quantitative marker overlap and lineage-consistency criteria. Functional summaries are generated deterministically from enrichment analysis of cluster marker genes. Specifically, GO-Slim enrichment is performed on the top-ranked marker genes using hyper-geometric testing with Benjamini–Hochberg correction (FDR ¡ 0.05). The top enriched terms are formatted into structured text and combined with the ranked marker list and filtered candidate labels to form the model input.

A small instruction-tuned language model is used without additional fine-tuning as a constrained annotator. For each cluster, the model is provided with (i) the top 20 marker genes, (ii) GO-Slim functional programs derived from enrichment analysis, and (iii) the highest-scoring gene program derived from the marker-based evidence module. The model is additionally supplied with the filtered candidate label set. Outputs are programmatically validated against this set. Responses not matching a valid label are returned as “None”.

#### Rationale for constrained language-model inference

A purely rule-based approach could select labels by maximizing marker overlap or enrichment statistics. However, enriched functional terms often reflect shared transcriptional programs (e.g., inflammation, stress response, extracellular matrix remodeling) rather than definitive cell-type identity, and multiple candidate labels may exhibit similar enrichment profiles. We therefore use a language model as a constrained semantic integrator over a curated candidate set, combining heterogeneous evidence (ranked markers, GO-Slim summaries, and label nomenclature/synonyms). This approach reduces reliance on brittle hand-crafted scoring rules while preventing unconstrained generation.

### 2.4 Confidence Scoring and Ambiguity Detection

For each cluster, candidate cell-type labels are scored by aggregating marker-gene evidence using the curated gene–label knowledge base. Each marker contributes a deterministic weight combining saturated log_2_ fold change, optional prevalence difference (pct_in_ − pct_out_), and adjusted *p*-value significance. Label scores are obtained by summing marker contributions across all supporting genes.

Marker coverage is defined as the fraction of top-ranked markers mapping to at least one candidate label. A normalized separation margin between the top two candidate scores quantifies label distinctiveness. Uncertainty is measured using Shannon entropy computed over softmax-normalized label scores.

A scalar confidence score integrates evidence strength, separation margin, and coverage, with entropy-dependent down-weighting. Default ambiguity thresholds were: minimum coverage 0.25, minimum top score 1.25, minimum margin 0.02, and maximum entropy 2.25.

Clusters failing any separability criterion are marked *AMBIGUOUS* and excluded from definitive labeling and quantitative evaluation. Otherwise, the top-scoring label is assigned and stratified by confidence.

### 2.5 Multi-Stage Refinement and Confidence-Based Abstention

Clusters identified as ambiguous are subjected to a second-stage refinement. For each such cluster with cell index set ℐ, we construct the induced subgraph of the original hybrid adjacency matrix, *A*^(2)^ = *A*_mix_[*I, I*] corresponding to the cells within that cluster. Leiden clustering is applied to *A*^(2)^ to identify potential subclusters.

Proposed splits are accepted only if they satisfy structural constraints, including minimum subcluster size (e.g., ≥200 cells or ≥5% of the parent cluster) and an upper bound on the number of subclusters. Accepted subclusters are re-annotated using the same constrained semantic procedure and confidence scoring described above.

If refinement does not produce subclusters meeting separability criteria, the framework abstains from assigning a definitive cell-type label. Such clusters are excluded from the final annotated label set and from quantitative evaluation. This design prevents overconfident labeling in biologically mixed or transitional regions.

### 2.6 Robustness, Ablation, and Evaluation

Robustness is assessed using a single pre-trained instruction-tuned model without additional fine-tuning and assess robustness via spatial ablation, performing clustering on the feature graph *A*_feature_ alone. Ambiguity rates and refinement outcomes are compared to the hybrid graph setting.

Performance is evaluated using top-*k* label agreement, confidence distributions, and the proportion of clusters requiring abstention. For clusters subjected to refinement, we additionally measure changes in confidence relative to initial assignments.

## 3 Experiments

### 3.1 Dataset and Implementation Details

We evaluate the proposed framework on 10x Genomics Xenium spatial transcriptomics data from cholangiocarcinoma. Gene expression matrices were normalized and clustered using the hybrid spatial–feature graph described in Section 2.1. Differential expression was computed using `scanpy.tl.rank_genes_groups`. Candidate labels were constructed from curated PanglaoDB and CellMarker references.Constrained semantic annotation was performed using `Qwen2.5-1.5B-Instruct`without additional fine-tuning. Unless otherwise stated, default ambiguity thresholds were: minimum coverage 0.25, minimum top score 1.25, minimum margin 0.02, and maximum entropy 2.25.

#### Clustering Configurations

We evaluate three clustering configurations. Hybrid 1-Stage performs single-stage Leiden clustering on the hybrid spatial–feature graph followed by constrained semantic annotation. Hybrid 2-Stage applies the same hybrid graph but introduces selective refinement of low-confidence clusters via local reclustering prior to re-annotation. Feature (2-Stage) follows the same two-stage refinement strategy as Hybrid 2-Stage but replaces the hybrid graph with the expression-based feature graph alone, excluding spatial adjacency.

### 3.2 Multi-Stage Refinement and Ambiguity Reduction

Table 1 summarizes the effect of multi-stage refinement and spatial integration. Compared to the single-stage hybrid pipeline, the two-stage refinement substantially reduces ambiguity at both cluster and cell levels, decreasing the proportion of ambiguous clusters from 16.1% to 2.27% and the fraction of affected cells from 18.4% to 0.86%. This reduction is accompanied by controlled sub-cluster discovery (23 to 44 clusters), indicating improved structural separability. Mean confidence increases modestly under refinement, reflecting improved evidence concentration within resolved clusters.

**Table 1:** Effect of multi-stage refinement and spatial integration. Amb. = ambiguous rate; Conf. = mean confidence (computed over non-ambiguous clusters).

| Method | Clusters |  |  | Cells |  |  |
| --- | --- | --- | --- | --- | --- | --- |
|  | N | Amb. (%) | Conf. | N | Amb. (%) | Conf. |
| Hybrid 1-Stage | 23 | 16.1 | 0.733 | 191125 | 18.4 | 0.719 |
| Hybrid 2-Stage | 44 | 2.27 | 0.744 | 191125 | 0.86 | 0.725 |
| Feature (2-Stage) | 28 | 7.14 | 0.766 | 191125 | 1.26 | 0.739 |

We further compare hybrid spatial–feature clustering against a feature-only baseline under the two-stage setting, where clustering is performed using the expression-based feature graph without incorporating spatial adjacency. Hybrid integration yields lower ambiguity rates at both cluster and cell levels, while maintaining comparable confidence calibration. Although the feature-only model exhibits slightly higher mean confidence, this corresponds to coarser partitioning rather than improved separability. Spatial integration improves structural resolution and reduces ambiguity without sacrificing reliability.

### 3.3 Qualitative Refinement Examples

We present representative clusters that were initially ambiguous at Hybrid 1-Stage and resolved through second-stage refinement. Table 2 summarizes the resulting cluster-level annotations, and spatial distributions before and after refinement are shown in Fig. 3.

**Table 2:**
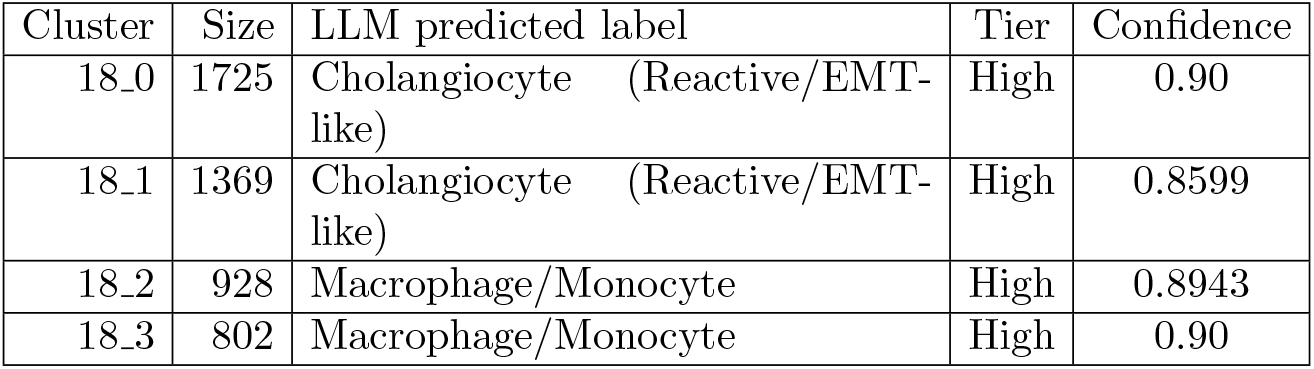
Cluster-level cell-type predictions with confidence tiers. The *Qwen label* denotes the raw LLM prediction, while the reported confidence reflects the strength and specificity of marker-gene support for the Qwen-predicted label after confidence-based ambiguity handling.

**Figure 1:**
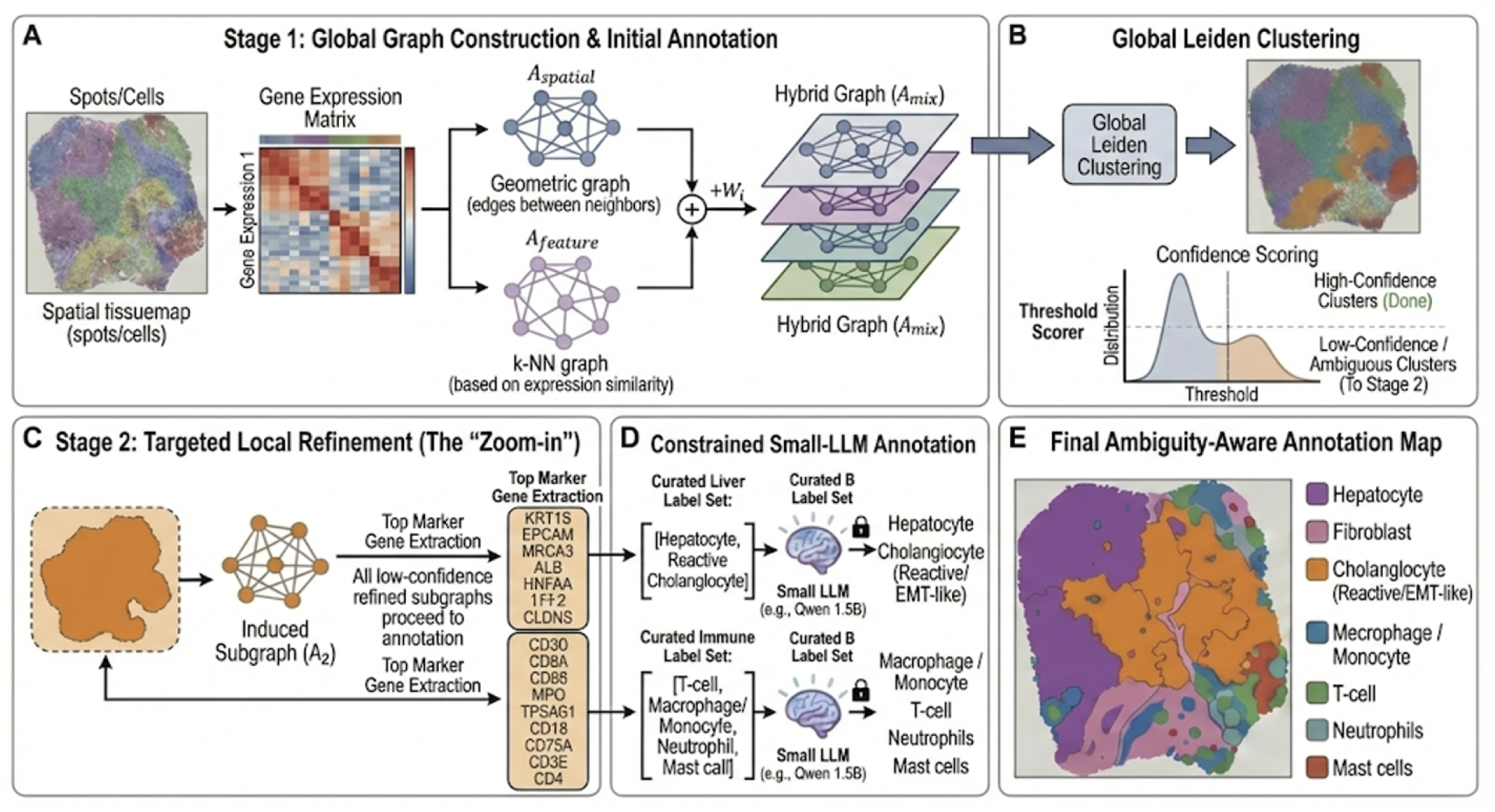
The ambiguity-aware multi-stage spatial annotation framework. (A) Stage 1 initiates with global graph construction, combining spatial coordinates and gene expression into a hybrid graph (*A*_*mix*_). (B) Global Leiden clustering is followed by confidence scoring, stratifying clusters into high-confidence (finalized) or low-confidence (ambiguous, requiring refinement) groups. (C) Stage 2 performs targeted local refinement (“Zoom-in”) on low-confidence subgraphs through marker gene extraction. (D) Constrained annotation utilizes curated label sets to guide a small LLM (e.g., Qwen 1.5B) in resolving cell identities. (E) The final output showing the refined, ambiguity-aware spatial annotation map.

**Figure 2:**
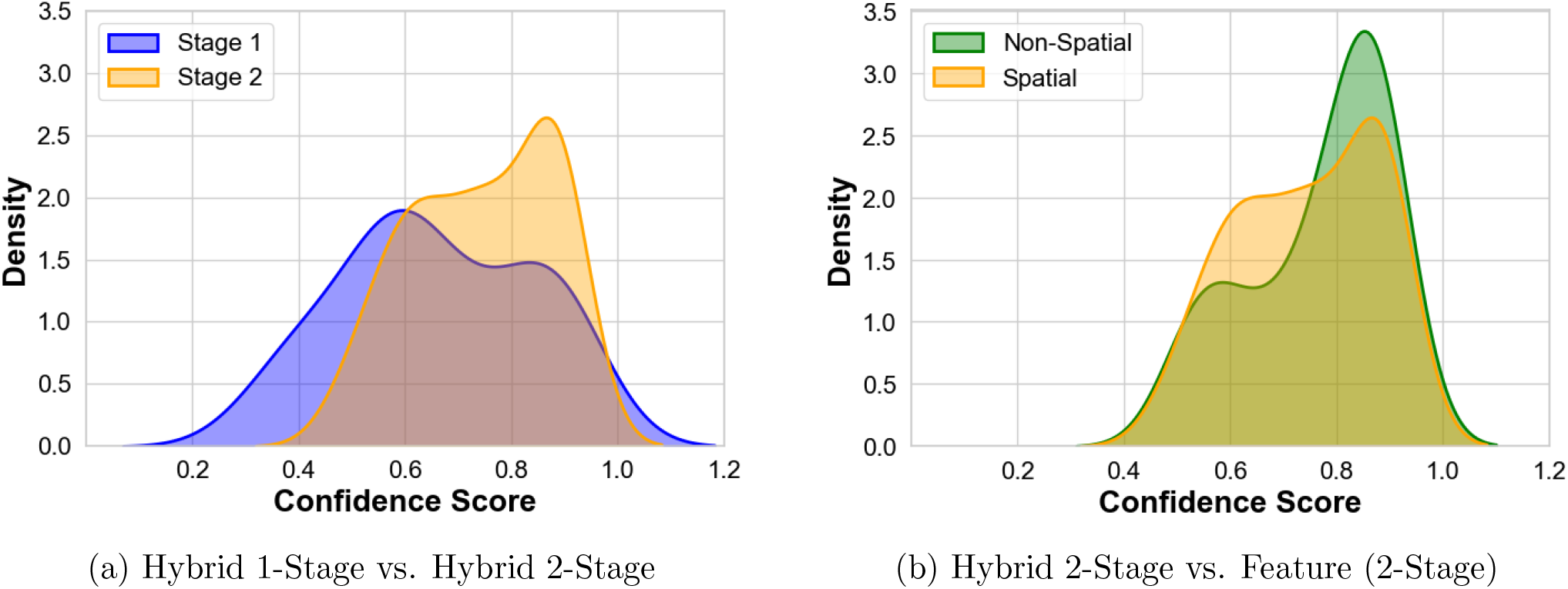
Confidence score density distributions. (a) Comparison of Hybrid 1-Stage and Hybrid 2-Stage annotations. (b)Comparison of our Hybrid 2-Stage framework against the Feature (2-Stage) baseline.

**Figure 3:**
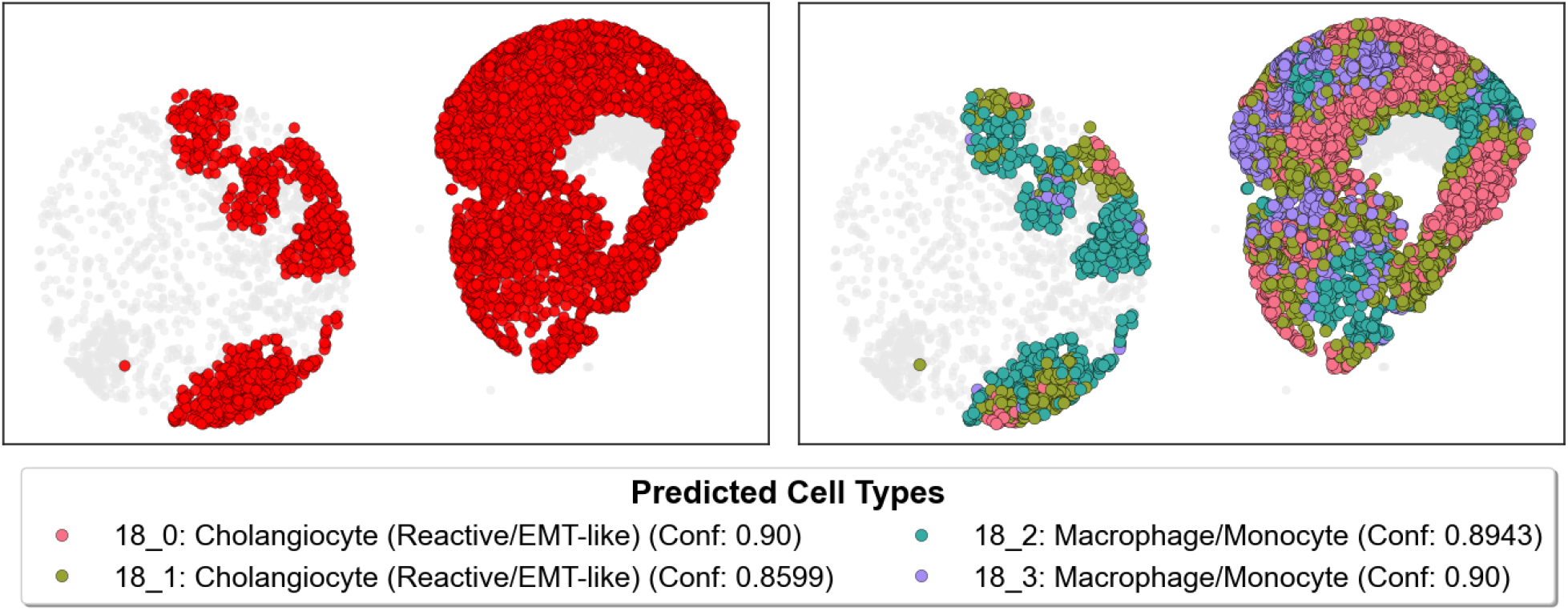
Core-level spatial resolution of ambiguous microenvironments. (Left) Initial global clustering (Hybrid 1-Stage) groups cells into a mixed, ambiguous region (Cluster 18, red) due to overlapping expression profiles. (Right) Our neighborhood-aware spatial refinement (Hybrid 2-Stage) successfully separates this mixed region into distinct, high-confidence cell populations. By leveraging local topology, the framework accurately isolates myeloid immune infiltrates (macrophages/monocytes) from reactive epithelial compartments (EMT-like cholangiocytes), demonstrating the critical role of spatial context in precise cell typing.

Refinement separates the ambiguous parent domain into two spatially and transcriptionally coherent populations: epithelial-like clusters (18 0, 18 1) and immune-like myeloid clusters (18 2, 18 3). The epithelial clusters are enriched for epithelial markers (e.g., *TACSTD2, EPCAM, SLPI*) together with inflammatory signaling genes (e.g., *CXCL8, CXCL1*), consistent with a reactive epithelial program. In contrast, clusters 18 2 and 18 3 exhibit immune-associated signatures including phagocytic and complement-related genes (e.g., *CD68, AIF1, C1QA/C1QC, MS4A7*), indicating myeloid identity.

Within each lineage, refinement further resolves state-level heterogeneity. The epithelial clusters differ in the relative strength of inflammatory and stress-response programs, while the myeloid clusters vary in lysosomal, complement, and inflammatory signaling components. These splits reflect structured transcriptional variation rather than noise.

Spatially, the two lineages occupy adjacent but distinct regions, indicating that the original ambiguous cluster represented a mixed epithelial–immune interface. Refinement therefore separates overlapping yet biologically distinct populations, yielding high-confidence annotations without imposing a single label on a heterogeneous domain.

## 4 Conclusion

We presented an ambiguity-aware, multi-stage framework for spatial cell-type annotation that integrates hybrid spatial–feature clustering, constrained language-model inference, and confidence-based abstention. By explicitly quantifying separability and selectively refining low-confidence clusters through induced subgraphs of the original hybrid graph, the method reduces ambiguous assignments while preserving structural resolution.

On cholangiocarcinoma spatial transcriptomics data, two-stage refinement decreased cluster-level ambiguity from 16.1% to 2.27% and cell-level ambiguity from 18.4% to 0.86%, while maintaining calibrated confidence estimates. Incorporating spatial adjacency further improved structural separability compared to feature-only clustering. Importantly, unresolved clusters are abstained rather than forcibly labeled, aligning annotation certainty with available molecular evidence.

Overall, these results demonstrate that explicit ambiguity handling and principled abstention provide a reliable and interpretable framework for spatial annotation in heterogeneous tumor tissues.

## 5 Acknowledgments

This project was supported by the National Center for Advancing Translational Sciences (NCATS), National Institutes of Health, through Grant Award Number UM1TR004539. The content is solely the responsibility of the authors and does not necessarily represent the official views of the NIH.

